# Investigating the Effects of Synbiotic Intervention on Working Memory, Attention, and Inhibitory Control in Healthy Young Women

**DOI:** 10.1101/2024.10.08.617166

**Authors:** Yeganeh Salimi, Behnaz Namdarzadeh, Fatemeh Dehghani Aarani, Abdolhossien Vahabi, Ehsan Rezayat

**Affiliations:** Department of Cognitive Sciences, Faculty of Psychology and Education, University of Tehran, Tehran, Iran; Department of Psychology, Faculty of Psychology and Education, University of Tehran, Tehran, Iran; School of Electrical and Computer Engineering, College of Engineering, University of Tehran, Tehran, Iran; School of Cognitive Sciences, Institute for Research in Fundamental Sciences (IPM), Niavaran, Tehran, Iran

**Keywords:** Synbiotic, Working Memory, Attention, Inhibitory Control, Gut-Brain Axis

## Abstract

The gut-brain axis plays a key role in the bidirectional communication between the gastrointestinal system and the brain, with gut microbiota significantly influencing cognitive function. While synbiotic and probiotic interventions have shown potential cognitive benefits, results across studies remain mixed. This randomized, controlled, repeated-measures study investigated the effects of a 15-day synbiotic intervention on cognitive performance in 28 healthy women (aged 18–32), assigned to either a synbiotic or no-treatment control group. Cognitive performance was assessed before and after the intervention using the Parametric Auditory Working Memory Task and the Integrated Visual and Auditory Continuous Performance Test (IVA-CPT). Compared to the control group, participants in the synbiotic group demonstrated selective improvements in attention, particularly in overall visual attention and visual sustained attention, as indicated by significant group-by-time interactions (ps < .05). No significant changes were observed in auditory working memory or inhibitory control. These results suggest that short-term synbiotic supplementation may selectively enhance visual attention without broadly affecting executive functions.

## Introduction

The gut-brain axis (GBA) is a complex communication network linking the gastrointestinal system and the brain, crucial for maintaining both mental and gut health. It regulates cognitive functions, and emotional states while also ensuring gastrointestinal homeostasis (Carabotti et al., 2015; Cryan et al., 2019; Eastwood et al., 2021). Central to this axis is the gut microbiota, a diverse community of microorganisms that influences immune system development, nutrient metabolism, and the production of essential biomolecules (Cryan et al., 2019; Oriach et al., 2016; Sarkar et al., 2018).

The symbiotic relationship between the host and gut microbiota plays a crucial role in shaping brain function and behavior through intricate signaling pathways. Accumulating evidence highlights that modulation of gut microbiota can profoundly impact cognitive processes (Cryan et al., 2019; Eastwood et al., 2021; Morais et al., 2021). Notably, distinct microbiota profiles have been associated with working memory performance, and furthermore, higher microbial diversity correlates with enhanced connectivity within brain networks underpinning working memory (Arnoriaga-Rodriguez et al., 2020; Cai et al., 2021). Beyond working memory, transferring ADHD-associated microbiota into germ-free mice impaired white/gray matter integrity and reduced connectivity in attention-related brain regions (Tengeler et al., 2020).

Furthermore, in humans, higher levels of microbial metabolites like butyrate correlated with stronger neural markers of response inhibition (Willis et al., 2025). These findings support the notion that gut-derived metabolites and the interaction between microbiota and the host can modulate neural mechanisms underpinning core executive functions.

Both probiotics, which provide health benefits when consumed in adequate amounts, and prebiotics, non-viable components that support beneficial microbes, have shown promising effects on working memory in preclinical studies (Coletto et al., 2022; Griffin et al., 2023; Li et al., 2022; Ohsawa et al., 2015; Yin et al., 2024), and on working memory, attention, and inhibitory response in clinical studies (Bloemendaal et al., 2021; Önning et al., 2023; Papalini et al., 2019; Serra et al., 2019; Adikari et al., 2020; Cardona et al.,2021; Elhossiny et al., 2023; Azuma et al., 2023; Shi et al., 2023; Trezzi et al., 2025). Since both probiotics and prebiotics target the gut, it’s clear that combining them can enhance their individual benefits and potentially amplify their effects on intestinal health. This combination, known as a synbiotic, is a mixture of live microorganisms and specialized substrates that are utilized by these microbes to deliver health benefits to the host (Ouwehand et al., 2007; Swanson et al., 2020).

Recent findings suggest that synbiotic supplementation may enhance key cognitive functions, particularly working memory and attentional control. Studies in animal models (Cruz-Martínez et al., 2024; Parois et al., 2021) and children (Nieto-Ruiz et al., 2022) have reported significant improvements in working memory following synbiotic intake. Moreover, in individuals with ADHD, synbiotic supplementation has been associated with improvements in attentional performance and inhibitory control (Trezzi et al., 2025).

A balanced gut microbiota is crucial for protecting the host from harmful microorganisms and maintaining overall homeostasis (Swanson et al., 2020). However, factors like antibiotic treatments, stress, Western diets, and environmental pollution can disrupt this balance (Leong et al,.2018; Cryan et al., 2019; Kendig et al., 2021). Such disruptions, common even in healthy individuals, can lead to gut dysbiosis, contributing to inflammatory, pathogenic, and metabolic disorders that impair executive functions, including working memory, attention and inhibitory control (Cryan et al., 2019; Patangia et al., 2022; Zhao et al., 2022; Czajeczny et al., 2023).

Therefore, modulating gut microbiota is important for enhancing cognitive function in both individuals with health conditions and the broader population.

This study investigates the cognitive impact of a synbiotic formulation combining *Lactobacillus* and *Bifidobacterium* strains with fructooligosaccharides (FOS), delivering 10LJ CFU per capsule. Using standardized neuropsychological assessments, we examine whether this specific blend can enhance key domains of executive functioning—namely, working memory, attention, and inhibitory control—in healthy young adult females. To the best of our knowledge, no prior research has evaluated the cognitive effects of this precise synbiotic combination and dosage in this population. We therefore hypothesize that this novel intervention will lead to significant improvements in executive performance, highlighting the potential of gut microbiota modulation as a pathway to cognitive enhancement.

## Materials and methods

Forty-eight female participants were initially recruited for this investigation. Three individuals voluntarily withdrew from the experiment, and seventeen did not meet the inclusion criteria.

Participants were excluded if they had acute or chronic illnesses, adhered to specific dietary plans, took any medications, had a body mass index (BMI) outside the normal range, consumed alcohol, had a possibility of pregnancy, suffered from immunodeficiency diseases, color blindness, vision or hearing impairments, or engaged in substance abuse. Additionally, individuals who had used probiotics, synbiotics, or antibiotics within the two months prior to the study were excluded. These criteria ensured the suitability of volunteers and minimized confounding variables that could affect the outcomes related to gut microbiota manipulation and cognitive performance. Considering the influence of gender on gut microbiota variation, all participants were female, aged between 18 and 32 years.

The study protocol was reviewed and approved by the Research Ethics Committee of the Faculty of Psychology and Education at the University of Tehran (Approval ID: IR.UT.PSYEDU.REC.1403.041), ensuring adherence to ethical standards. Each participant provided written informed consent and was informed of their right to withdraw from the study at any time.

We conducted a repeated-measures design study including a control group (Figure 1). Initially, 28 participants were screened for depression and anxiety using the Beck Depression Inventory (BDI) and the Beck Anxiety Inventory (BAI). Demographic information was also collected at this stage. In the first session, participants completed the Positive and Negative Affect Schedule (PANAS), followed by a parametric auditory working memory task (Akrami et al,. 2018) and the Integrated Visual and Auditory Continuous Performance Test (IVA-2). After assessments, participants were randomly assigned to either the intervention (n=14) or control (n=14) groups.

**Figure 1:**
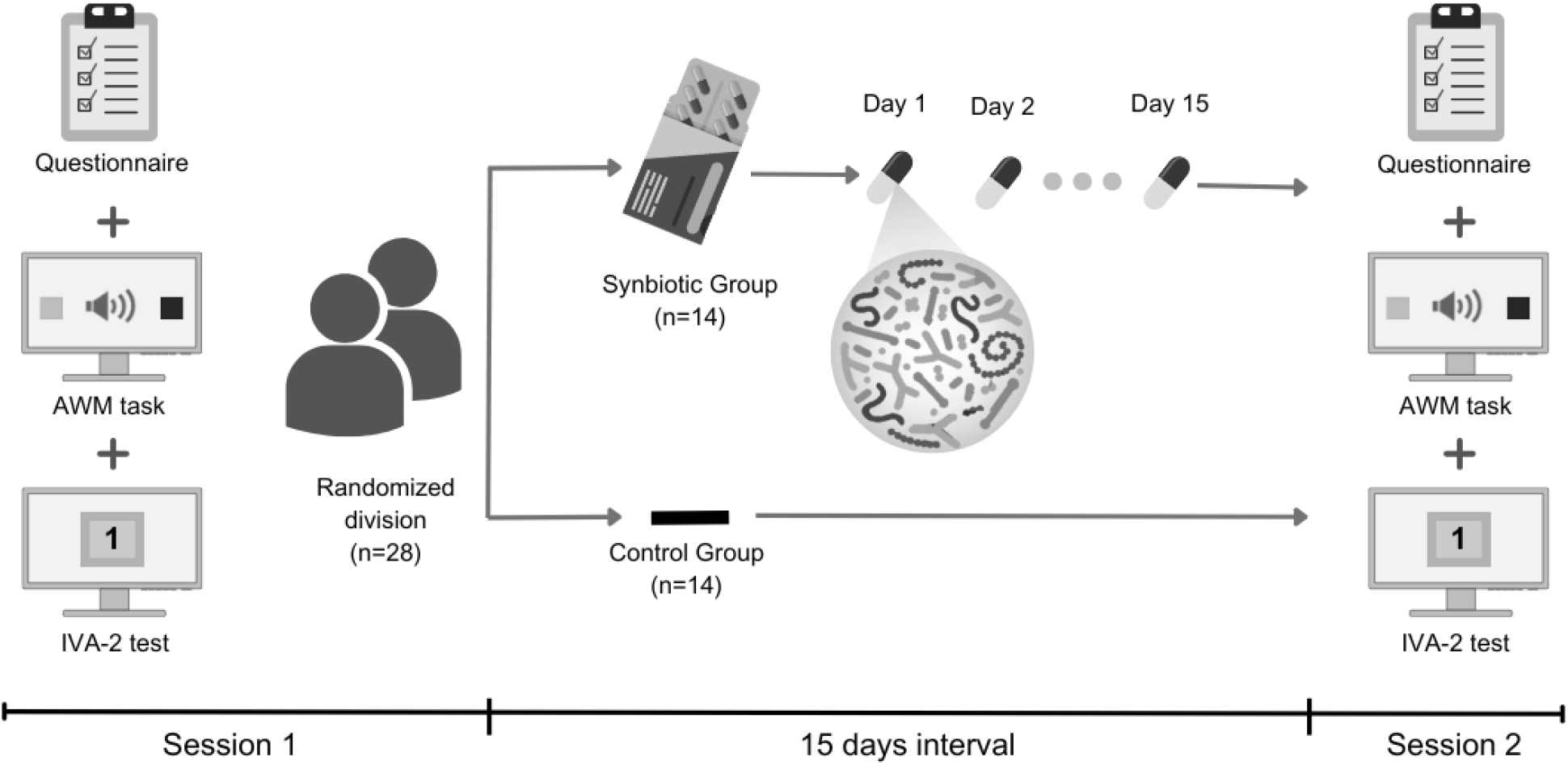
Experimental Design. The figure illustrates the repeated measures design, including the main procedures for both experimental sessions. It outlines the screening, random assignment to the intervention or control group, and the synbiotic capsule protocol followed by participants over 15 days.

Participants in the intervention group began taking synbiotic capsules, instructed to consume one capsule daily for 15 days. Adherence was closely monitored; participants missing two or more doses were excluded. Both groups were asked to maintain their usual diet and exercise routines and avoid other probiotics. Participants were also required to inform researchers of any antibiotic or medication use during the study, which could lead to exclusion. Intervention participants were advised to discontinue capsule use and contact researchers immediately if side effects occurred.

In the second session, all participants repeated the PANAS questionnaire, the auditory working memory task, and the IVA-2 test.

### Positive and Negative Affect Schedule

The PANAS questionnaire (Watson et al., 1988) is a self-report instrument designed to measure positive and negative affect. It consists of a series of adjectives describing various emotions.

Participants are asked to rate the intensity with which they have experienced each emotion over a specified time period, using a scale from 1 (very slightly or not at all) to 5 (extremely). Higher scores on the Positive Affect (PA) and Negative Affect (NA) scales indicate greater levels of positive and negative affect, respectively. In this study, participants completed the PANAS questionnaire online via Porsline.com at the beginning of each session, responding based on their mood during the preceding week.

### Parametric auditory working memory test

To assess auditory working memory performance, participants were presented with pairs of pink noise sounds at one of the following levels: 60, 62, 64, 66, 68, 70, or 72 dB, delivered through noise-cancelling headphones. The pairs of sounds were randomly assigned. Participants began the task by reviewing the instructions and completing six training trials. During the testing phase, the first auditory stimulus was presented simultaneously with the appearance of a green square on the left side of a computer monitor placed in front of the participant. After a delay period of either 2 or 6 seconds, indicated by the word “WAIT!” displayed on the screen, the second sound was presented along with a red square appearing on the right side of the screen. Following the second stimulus and the “go” cue, participants judged which of the two sounds was louder by clicking on the corresponding square using a mouse. Each participant completed a total of 150 trials (Figure 2A). The task was programmed in MATLAB (The MathWorks Inc., Massachusetts, US) using Psychtoolbox.

**Figure 2:**
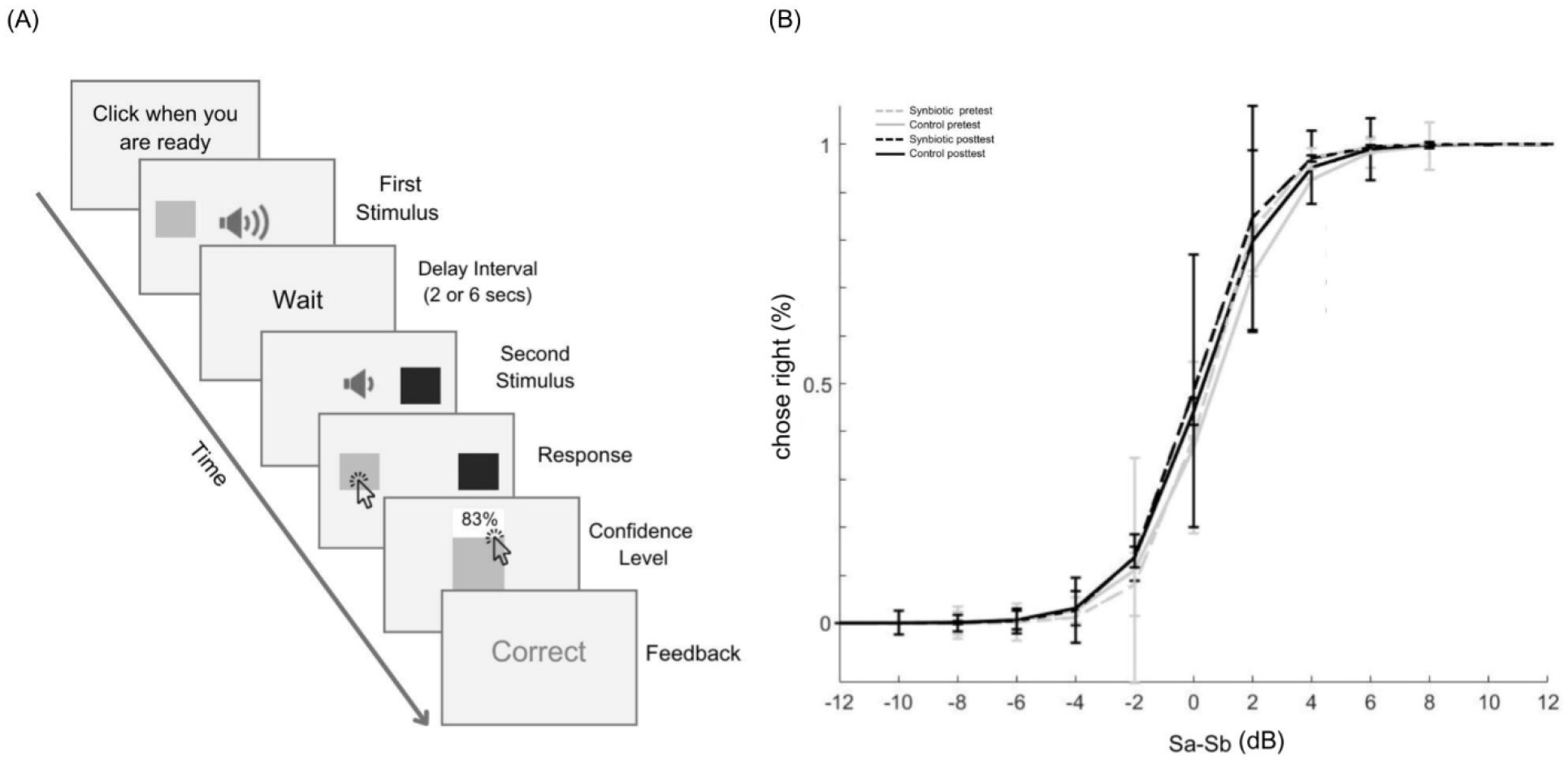
Parametric Auditory Working Memory. **A**, The figure shows the auditory delayed comparison task. Participants compared the loudness of two pink noise sounds, presented with a delay and marked by colored squares. They indicated their choice and confidence, with feedback provided after each of the trials. **B**, The figure shows GLM-fitted psychometric curves for both groups, depicting the probability of choosing the second stimulus (rightward) based on the dB difference between the first and second stimuli. The y-axis represents the percentage of rightward choices, and the x-axis indicates the stimuli difference.

### IVA-2

To assess attentional functioning and inhibitory control, we used the second edition of the Integrated Visual and Auditory Continuous Performance Test (IVA-2 CPT), developed by BrainTrain Inc. (USA). A Persian version of this computerized test, commonly used in research settings in Iran, was administered. During the task, participants sat approximately 40–60 cm from a computer monitor, with a two-button mouse placed in front of them. They were exposed to a total of 500 rapidly presented stimuli, consisting of both visual and auditory presentations of the numbers “1” and “2.” Each stimulus was displayed for 500 milliseconds. Participants were instructed to click the mouse when they saw or heard the number “1” (target stimulus) and to withhold their response when presented with “2” (non-target stimulus). The test was composed of three sequential phases: a brief warm-up, the main task (which included both practice and test segments), and a cool-down period. The entire procedure took approximately 20 minutes to complete. The IVA-2 generates eight global composite quotient scores and nineteen additional scales. These indices provide a multidimensional profile of attention and executive functioning across both auditory and visual modalities. In our analyses, we examined a comprehensive set of these outcomes—including global scales, modality-specific scores, and other cognitive– behavioral metrics.

### Supplementation

Synbiotic intake was controlled by administering a commercially available supplement (Familact 2Plus, Zist Takhmir, Iran) containing live bacterial species, including *Lactobacillus acidophilus, Lactobacillus casei, Lactobacillus rhamnosus, Lactobacillus salivarius, Lactobacillus reuteri, Bifidobacterium lactis, Bifidobacterium longum*, and *Bifidobacterium bifidum*, along with FOS as a prebiotic (10^9 CFU per capsule). The supplement was provided in enteric-coated capsules. Participants in the intervention group were instructed to follow the information sheet guidelines, taking one capsule on an empty stomach, 1 hour before a meal, for 15 days. No participants reported any side effects during or after the study.

### Statistical Analysis

A mixed factorial design was employed, with group (synbiotic vs. control) as the between-subjects factor and time (pretest vs. posttest) as the within-subjects factor. All statistical analyses were conducted in MATLAB, with the significance threshold set at p < 0.05.

Baseline variables—including age, BMI, BAI, BDI, and pretest PANAS scores—were compared between groups using independent-samples t-tests. Due to a violation of normality, years of education were analyzed using the non-parametric Mann–Whitney U test.

To examine intervention effects over time and between groups, linear mixed-effects models (LMEs) were applied. Each model included fixed effects for group, time, and their interaction (group × time), with random intercepts for participants to account for repeated measures. The same model structure was used to analyze PANAS and IVA-2 assessment scores.

Performance on the Auditory Working Memory (AWM) task was modeled at the individual level using binomial generalized linear models with a logit link, estimating psychometric parameters (intercept, slope, threshold, and sensitivity) for each participant across conditions and timepoints (Figure 2B). Subsequently, threshold, sensitivity, and log-transformed response times were analyzed using LMEs. Trial-level accuracy (binary) was analyzed using generalized linear mixed-effects models with identical fixed and random effects structures.

## Results

We included 28 female participants in the experiment, randomly assigned to either the control group (n = 14) or the synbiotic group (n = 14). Table 1 presents the demographic and baseline characteristics of the participants. No significant differences were observed between the two groups in terms of age, BMI, years of education, BAI, BDI, or baseline PANAS scores (*p* > .05 for all comparisons).

**Table 1:** the demographic and baseline characteristics of the participants.

| characteristics | Synbiotic group | Cotrol group | P-value |
| --- | --- | --- | --- |
|  | Mean(SD) | Mean(SD) |  |
| Age | 25.21(3.14) | 23.07(3.2) | 0.08 |
| BMI | 24.44(3.74) | 23.45(4.25) | 0.52 |
| Years of Education | 18.29(1.73) | 17.14(1.03) | 0.053 |
| BAI | 8.57(4.35) | 8.71(4.18) | 0.93 |
| BDI | 8.21(3.42) | 8.21(4.06) | 1 |
| PANAS Positive (pretest) | 29.43 (8.27) | 31.50 (8.22) | 0.51 |
| PANAS Negative (pretest) | 27.43 (7.98) | 24.57 (6.55) | 0.30 |

### PANAS score

Linear mixed-effects models were used to evaluate the effects of the synbiotic intervention on both positive and negative affect scores over time. For positive affect, no significant group × time interaction was observed (β = -0.07,SE= 2.8773, *p* = 0.98).

For negative affect, the group × time interaction was marginally significant (β = -5.86, SE= 3.0368, *p* = 0.059), with a small effect size (partial R^2^ = 0.0605), suggesting a modest proportion of variance explained by the group × time interaction.

### Auditory Working Memory Performance

For the threshold variable, a significant main effect of time was observed (β = -0.41, SE = 0.17, *p* = 0.020), indicating overall improvement across both groups over time. No significant effects of group (*p* = 0.46) or group × time interaction (β = -0.10, SE = 0.24, *p* = 0.69) were found (Figure 3).

**Figure 3:**
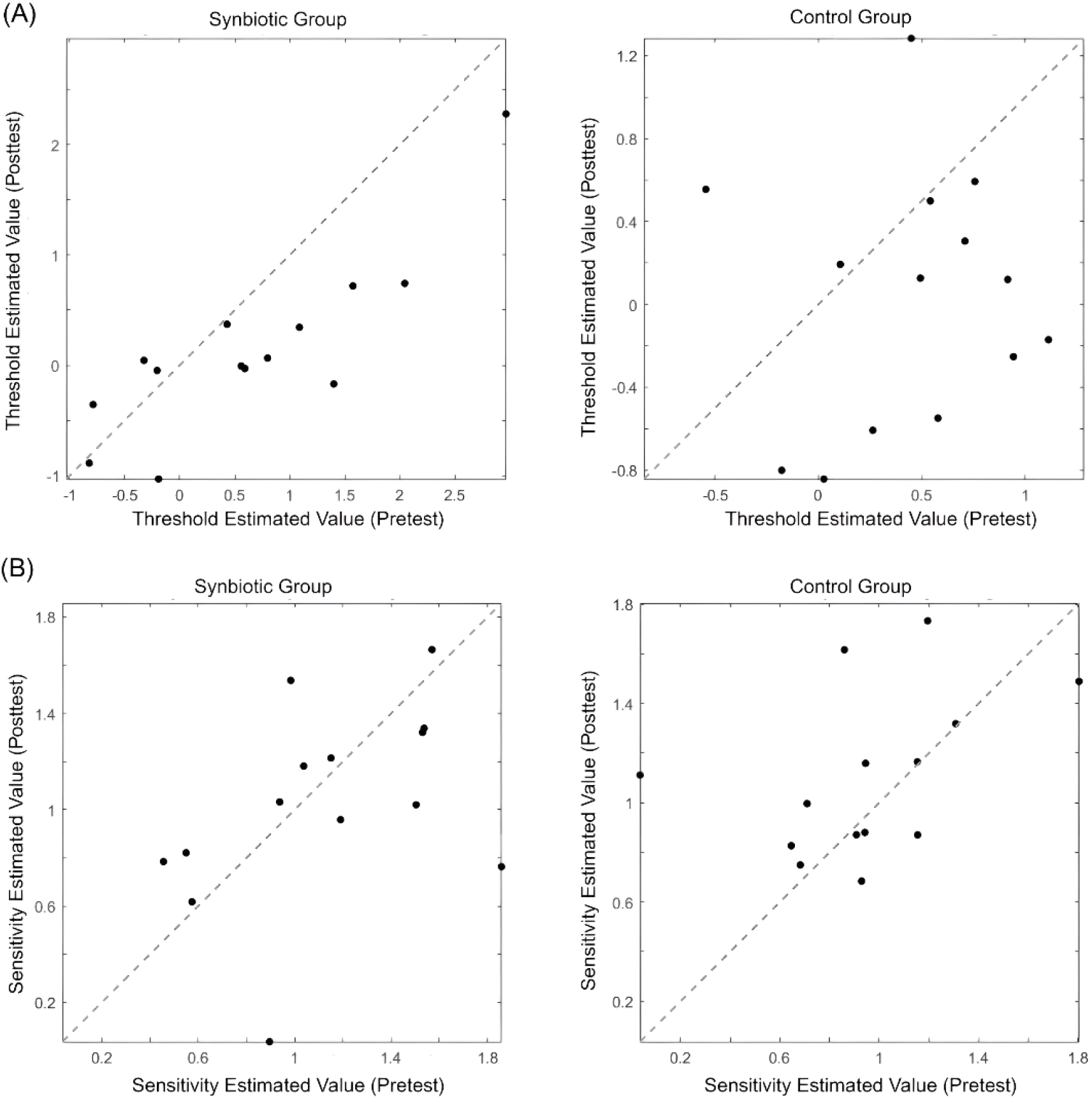
Threshold & Sensitivity. Scatter plots of pretest and posttest scores are shown for both the synbiotic group (left) and control group (right). Panel **A** depicts auditory threshold scores, while Panel **B** illustrates sensitivity scores. Each point represents an individual score, with the solid line indicating equality (y = x).

**Figure 4:**
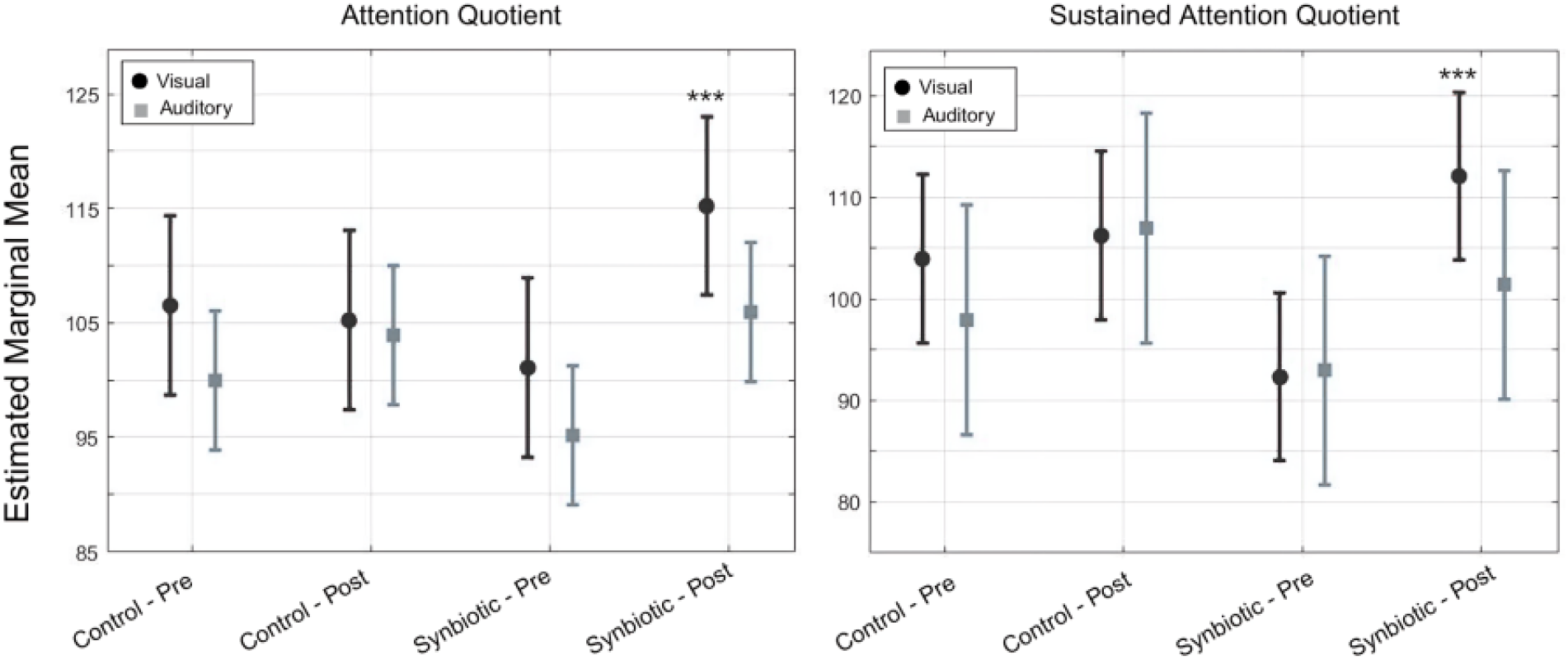
Estimated marginal means of Attention Quotient (left) and Sustained Attention Quotient (right) across groups (Control vs. Synbiotic) and sessions (Pre vs. Post), separated by modality (Visual vs. Auditory). A significant Group × Session interaction was observed for the Visual Attention Quotient *p* = .0244, and for the Visual Sustained Attention Quotient (*p* = .0212. These findings indicate a significant post-intervention improvement in the visual modality within the synbiotic group (\*\**p* < .001). Error bars represent 95% confidence intervals.

Regarding sensitivity, none of the fixed effects reached significance, although the group × time interaction showed a trend toward significance (β = -0.26, SE = 0.16, *p* = 0.098), with a small effect size (partial R^2^ = 0.0473) (Figure 3).

No significant main or interaction effects were found for group, time, or their interaction (all *p* > 0.08) in accuracy.

For reaction time (logRT), a significant main effect of time was observed (β = –0.08, SE = 0.014, p < .001), indicating faster responses over time across both groups. A significant main effect of group (β = –0.23, SE = 0.081, p = .004) also indicated that, overall, participants in the intervention group responded faster than those in the control group. However, a significant group × time interaction (β = +0.20, SE = 0.019, p < .001) revealed that the reduction in reaction time from pre-to post-test was greater in the control group than in the synbiotic group. The partial R^2^ for the reaction time interaction was small but positive (partial R^2^ = 0.0147).

### Attention

A significant group-by-time interaction was observed for the Full-Scale Attention Quotient (FSAQ), β = 13.14, SE = 5.26, p = .0156, with a partial R^2^ of 0.1008, indicating a greater improvement in the intervention group compared to the control group. Similarly, the Visual Attention Quotient (VAQ) demonstrated a significant interaction effect, β = 15.43, SE = 6.66, p= .0244, partial R^2^ = 0.0877. The Visual Sustained Attention Quotient (SAVQ) also showed a significant group-by-time interaction, β = 17.50, SE = 7.37, p = .0212, partial R^2^ = 0.0918, suggesting a notable post-intervention improvement in the intervention group (Figure 1). A significant interaction was found for Visual Focus, β = 13.21, SE = 6.24, p = .0390, partial R^2^ = 0.0743. Similarly, Combined Mental Concentration exhibited a significant interaction, β = 8.93, SE = 4.32, p = .0436, partial R^2^ = 0.0711. The strongest effect was observed for Visual Mental Concentration, with a substantial group-by-time interaction, β = 19.79, SE = 6.39, p = .0031, partial R^2^ = 0.1473, highlighting the intervention’s pronounced effect on focused attention.

No other significant main effects were found (ps > .05). Descriptive statistics and full model results are provided in Supplementary Table 1.

### Complementary Cognitive Abilities

Results revealed a significant main effect of group on visual presence (β = -9.86, SE = 4.55, *p* = 0.035), as well as a significant group × time interaction (β = 13.86, SE = 5.97, *p* = 0.024, partial R^2^ =0.088), indicating improved performance in the intervention group post-treatment.

Similarly, for the Steadiness index, a significant negative main effect of group was observed (β = -17.64, SE = 4.94, *p* = 0.00077), alongside a significant positive group × time interaction (β = 18.36, SE = 6.77, *p* = 0.009, partial R^2^ = 0.1182), reflecting enhanced steadiness following the intervention. The visual Steadiness measure showed comparable findings, with a significant negative group effect (β = -15.21, SE = 6.66, *p* = 0.026) and a positive group × time interaction (β = 19.71, SE = 8.23, *p* = 0.020, partial R^2^ = 0.0913), suggesting improved steadiness over time in the intervention group.

Furthermore, indices of visual Resilience and visual Stability demonstrated significant positive group × time interactions (β = 17.79, SE = 8.56, *p* = 0.043, partial R^2^ = 0.0717 ; β = 13.00, SE = 5.49, *p* = 0.022, partial R^2^ =0.0913, respectively), indicating beneficial effects of the probiotic intervention on cognitive resilience and stability. Other outcome measures did not exhibit significant group or interaction effects. Comprehensive results for these measures are provided in the supplementary Table S1.

### IVA-2 Symptomatic Scales

Among the symptom-based measures, significant group × time interactions were found for Visual Comprehension and Visual Sensorimotor Functioning. In Visual Comprehension, the interaction was significant (β = 9.93, SE = 3.80, *p* = .012, partial R^2^ = 0.1090), indicating a differential change over time between groups. There was also a significant group main effect (β= –9.43, SE = 4.35, *p* = .035), with the intervention group showing lower baseline scores. Similarly, for Visual Sensorimotor Functioning, the group × time interaction was significant (β = 5.00, SE = 2.24, *p* = .030, partial R^2^ = 0.0818), along with a significant group main effect (β = – 5.57, SE = 2.73, *p* = .046). None of the other symptomatology subscales showed significant effects; full model details are reported in Supplementary Table S1.

### Inhibitory Control

None of the models examining inhibitory control scales revealed any statistically significant main or interaction effects. For Full Scale Response Control Quotient (FSRCQ), the group × time interaction was not significant (*p* = .79). Likewise, no significant interaction effects were observed for Prudence (*p* = .9), Consistency (*p* = .53), Stamina (*p* = .5), Hyperactive events (*p* = .31), Fine motor hyperactivity (*p* = .48), and Self-control composite (*p* = .51). Complete model outputs, including auditory/visual subscores, are reported in Supplementary Table 1.

## Discussion

This study investigated the effects of a short-term synbiotic intervention on executive functions in healthy young women. While no significant changes were observed in auditory working memory or inhibitory control, the synbiotic group demonstrated greater improvements in visual attention compared to controls. These findings suggest that synbiotic supplementation may selectively enhance visual attentional performance.

Baseline comparisons confirmed that the synbiotic and control groups were well matched across key demographic and psychological variables—including age, BMI, education, anxiety, depression, and baseline mood—enhancing confidence that observed cognitive changes are attributable to the intervention rather than pre-existing differences.

Although no meaningful changes emerged in positive affect, a marginal trend toward reduced negative affect was observed in the synbiotic group, suggesting a potential subtle influence on mood-related processes. Given the small effect size and borderline significance, this finding should be interpreted with caution and warrants further investigation in larger, longer-term trials.

The existing literature on probiotics and mood presents mixed results, consistent with our findings. For example, Bagga et al. (2018) reported significant improvements in positive affect using a probiotic blend that included strains also used in our study (e.g., *B. bifidum, B. lactis, B. longum, L. acidophilus, L. casei, L. salivarius*). In contrast, Boehme et al. (2023) found that *B. longum* alone produced only a modest increase in positive affect and no change in negative affect. These discrepancies—both in prior studies and in our own data—underscore the complex and strain-dependent nature of probiotic effects on mood.

Both the synbiotic and control groups showed significant reductions in auditory thresholds over time. However, the absence of between-group differences suggests that improvements in auditory detection are likely attributable to practice effects rather than the synbiotic intervention. While there was a non-significant trend toward group differences in auditory sensitivity, the small effect size indicates that synbiotic supplementation did not meaningfully alter auditory discrimination in this study.

Similarly, no significant effects were found on auditory working memory accuracy. Although performance improved modestly over time, neither synbiotic intake nor repeated testing appeared to exert a meaningful influence. Analysis of reaction times revealed that only the control group showed significant reductions over time; in contrast, the synbiotic group exhibited minimal change—or in some cases, slight increases. This finding contrasts with studies suggesting that probiotics may improve processing speed, at least under certain conditions.

Our findings align with some prior research. For instance, Romo-Araiza et al. (2023) reported no significant differences in working memory accuracy between synbiotic and control groups in rats. Similarly, Moloney et al. (2021) observed no improvement in working memory following *Bifidobacterium longum* supplementation—one of the strains included in our formulation. On the other hand, studies such as Cruz-Martínez et al. (2024) reported enhanced working memory accuracy in ischemic stroke rats following a similar synbiotic combination, suggesting that such benefits may be population-dependent.

The lack of significant improvement in reaction times within the synbiotic group also diverges from some studies reporting enhanced processing speed. For example, Sanborn et al. (2018) found improved processing speed after *Lactobacillus GG* supplementation in middle-aged and older adults, while Adikari et al. (2020) observed faster reaction times in athletes following *Lactobacillus casei Shirota* intake. These inconsistencies highlight the complex interplay of multiple factors—including strain composition, prebiotic type, duration, dosage, and participant characteristics (e.g., age, health status, baseline cognition)—which may collectively determine whether synbiotic supplementation exerts measurable cognitive benefits.

The present findings indicate that synbiotic supplementation significantly enhanced attentional performance, particularly within the visual domain. Specifically, the intervention group demonstrated greater post-intervention gains in FSAQ, VAQ, VSAQ, Visual Focus, and Visual Mental Concentration compared to the control group. These indices reflect the ability to detect, focus on, and maintain consistent responses to visual stimuli over time. The corresponding effect sizes (partial R^2^ ranging from 0.0711 to 0.1473) suggest that these improvements were not only statistically significant but also of meaningful magnitude, particularly for visual mental concentration, which showed the largest effect.

Several complementary cognitive indices further supported the selective enhancement of visual attention following synbiotic supplementation. Notably, Visual Presence exhibited a significant group-by-time interaction, indicating improved readiness and engagement with visual stimuli after the intervention. Similarly, the Steadiness index showed significant improvements, suggesting increased consistency in attentional control. Moreover, Visual Resilience and Visual Stability were significantly enhanced in the synbiotic group, underscoring the intervention’s positive impact on cognitive stability and sustained visual attention. The effect sizes for these outcomes (partial R^2^ ranging from 0.0717 to 0.1182) suggest that these improvements were of small to moderate practical significance

Importantly, no significant improvements were observed in auditory-based attention indices. However, certain visual indices (e.g., Visual Agility, Visual Quickness, Visual Swiftness) approached significance, suggesting a potential trend toward improvement that did not reach conventional thresholds.

Our findings differ from those of Moloney et al. (2020), who reported no improvement in general attention following probiotic use. Conversely, they align with results from Elhossiny et al. (2023), who observed enhanced sustained attention in children with ADHD after probiotic supplementation, and Adikari et al. (2020), who found improved attentional performance under stress in healthy adults. These mixed findings in the literature highlight the importance of context, participant characteristics, and specific probiotic formulations in shaping cognitive outcomes.

In addition to performance-based improvements, synbiotic supplementation was associated with significant gains in Visual Comprehension and Visual Sensorimotor Functioning. These measures reflect the capacity to interpret visual information and coordinate visual input with motor responses in a goal-directed manner. The observed effect sizes indicate small to moderate magnitudes of change, suggesting that these improvements were not only statistically significant but also of practical relevance. In contrast, no significant changes were observed in auditory-based symptom indices. These results are consistent with prior research demonstrating that synbiotic and probiotic interventions can affect sensorimotor outcomes in neurological conditions such as stroke and traumatic brain injury (Rahman et al., 2024; Holcomb et al., 2025), supporting the potential role of gut microbiota modulation in sensory and motor functioning.

This selective enhancement pattern implies that the cognitive benefits of synbiotic supplementation may be modality-specific to visual attentional systems compared to auditory cognitive functions. From a functional perspective, such improvements may translate to enhanced stability, precision, and mental endurance in tasks requiring sustained visual engagement. Furthermore, no significant effects of synbiotic supplementation were observed on any measures of inhibitory control in healthy adult females. Both overall response control and its subcomponents—such as prudence, consistency, stamina, hyperactivity, fine motor control, and self-control—remained unchanged throughout the intervention period. These indices capture various aspects of response inhibition and impulse regulation, including the ability to withhold inappropriate responses, maintain consistent behavioral output, and manage impulsive or hyperactive tendencies. The absence of improvements suggests that synbiotic supplementation may not substantially influence the neural or behavioral mechanisms underlying inhibitory control in this context. These findings align with previous studies reporting minimal or no benefits of probiotic supplementation on executive and inhibitory functions in healthy populations (Edebol Carlman et al., 2022; Czajeczny et al., 2023), underscoring the need for further research to identify conditions under which gut–brain interventions might effectively modulate inhibitory control.

Several limitations must be considered. First, the research exclusively involved female participants, limiting the generalizability of the findings to males. Second, the study had a short intervention period, potentially overlooking the long-term benefits of synbiotics on cognitive function. Future research should explore different dosages of synbiotics and employ longer intervention durations. Moreover, examining changes in microbiota profiles post-intervention would provide valuable insights into the underlying mechanisms, ensuring that observed cognitive changes result from bacterial colonization in the gut rather than from factors like paraprobiotics. Finally, there is currently no direct evidence elucidating the specific mechanistic pathways through which gut microbiota might modulate visual attention. Most mechanistic studies to date have focused on broader cognitive functions or emotional aspects of attention (i.e., hot cognition), rather than the cold cognitive processes relevant here. This gap in the literature limits our ability to precisely interpret the pathways mediating the observed effects. By addressing these limitations, future studies could yield more robust evidence of the cognitive benefits of synbiotic supplementation.

