## Supplementary material for "Investigating the Effects of Synbiotic Intervention on Working Memory, Attention, and Inhibitory Control in Healthy Young Women": Supp

*Table 1 S1. includes all IVA measures where the linear mixed-effects models did not show statistically (group*time)*

| Scale | estimate | SE | tstat | p-value |
|---|---|---|---|---|
| Auditory Attention Quotient | 6.7857 | 5.4895 | 1.2361 | 0.22 |
| Sustained Attention Quotient | 8.9286 | 5.4521 | 1.6377 | 0.10753 |
| Sustained Auditory Attention Quotient | -0.57143 | 7.5469 | -0.075717 | 0.93994 |
| Vigilance | 3 | 4.6527 | 0.64478 | 0.5219 |
| Auditory Vigilance | 0.92857 | 5.7061 | 0.16273 | 0.87136 |
| Visual Vigilance | 9.7143 | 5.7795 | 1.6808 | 0.098799 |
| Focus | 6.0714 | 5.4721 | 1.1095 | 0.27231 |
| Auditory Focus | 5.5 | 5.4121 | 1.0162 | 0.31422 |
| Speed | 5.9286 | 4.3343 | 1.3678 | 0.17725 |
| Auditory Speed | 6.1429 | 6.9633 | 0.88218 | 0.38174 |
| Visual Speed | 5.2143 | 4.1603 | 1.2533 | 0.21568 |
| Auditory Mental Concentration | 1.3571 | 4.531 | 0.29953 | 0.76573 |
| Full Scale Response Control Quotient | 1.4286 | 5.426 | 0.26328 | 0.79337 |
| Auditory Response Control Quotient | 3.7857 | 4.7501 | 0.79698 | 0.42909 |
| Visual Response Control Quotient | 2.2857 | 7.0339 | 0.32496 | 0.74652 |
| Prudence | -1.0714 | 8.6641 | 0.12366 | 0.90206 |
| Auditory Prudence | 6.0714 | 4.977 | 1.2199 | 0.228 |
| Visual Prudence | -7.0714 | 11.39 | -0.62085 | 0.53741 |
| Consistency | 3.0714 | 4.8596 | 0.63204 | 0.53013 |
| Auditory Consistency | 7.7857 | 5.1855 | 1.5014 | 0.13929 |
| Visual Consistency | 7.5714 | 6.0428 | 1.253 | 0.21583 |
| Stamina | -3.5714 | 5.3754 | -0.6644 | 0.50937 |
| Auditory Stamina | -2.5714 | 6.9076 | -0.37226 | 0.71121 |
| Visual Stamina | 0.71429 | 5.8002 | 0.12315 | 0.90246 |
| Hyperactive Events | 1.6429 | 1.6213 | 1.0133 | 0.31561 |
| Fine Motor Hyperactivity | -4.1429 | 5.9464 | -0.6967 | 0.4891 |
| Self-Control | -6.0714 | 9.2742 | -0.65466 | 0.51557 |
| Auditory Self-Control | -8.2857 | 12.984 | -0.63813 | 0.52619 |
| Visual Self-Control | 3.0714 | 9.3326 | 0.32911 | 0.7434 |
| Presence | 7.9286 | 4.9783 | 1.5926 | 0.11731 |
| Auditory Presence | 1.4286 | 5.8373 | 0.24473 | 0.80763 |
| Acuity | 4.6429 | 5.0373 | 0.9217 | 0.36094 |
| Auditory Acuity | -1.2857 | 4.5136 | -0.28485 | 0.77689 |
| Visual Acuity | 9 | 6.5179 | 1.3808 | 0.17324 |
| Elasticity | -0.92857 | 4.3299 | -0.21445 | 0.83103 |
| Auditory Elasticity | -5.6429 | 6.0425 | -0.93387 | 0.35469 |
| Visual Elasticity | 2.0714 | 5.0871 | 0.40719 | 0.68554 |
| Auditory Steadiness | 15.214 | 8.0114 | 1.8991 | 0.0631 |
| Resilience | 10.786 | 5.9576 | 1.8104 | 0.076012 |
| Auditory Resilience | 2.7857 | 5.3594 | 0.51978 | 0.60542 |

| | | | | |
|---|---|---|---|---|
| Stability | 8.7143 | 5.4052 | 1.6122 | 0.11297 |
| Auditory Stability | 5.3571 | 5.4717 | 0.97907 | 0.33208 |
| Agility | 3.5714 | 4.7753 | 0.74789 | 0.4579 |
| Auditory Agility | 2.2857 | 5.3573 | 0.42666 | 0.67139 |
| Visual Agility | 8.7143 | 4.6523 | 1.8731 | 0.066681 |
| Quickness | 4.5 | 4.6154 | 0.97499 | 0.33408 |
| Auditory Quickness | 2.5714 | 5.6912 | 0.45182 | 0.65327 |
| Visual Quickness | 8.0714 | 4.2881 | 1.8823 | 0.065401 |
| Swiftness | 4.8571 | 4.8547 | 1.0005 | 0.3217 |
| Auditory Swiftness | 0.85714 | 5.2396 | 0.16359 | 0.87069 |
| Visual Swiftness | 8.5 | 4.8772 | 1.7428 | 0.087282 |
| Accuracy | 7.3571 | 4.3519 | 1.6906 | 0.096909 |
| Auditory Accuracy | 4.8571 | 3.7062 | 1.3105 | 0.19577 |
| Visual Accuracy | 5.8571 | 5.3005 | 1.105 | 0.27424 |
| Auditory Comprehension | 5.5 | 3.5395 | 1.5539 | 0.12628 |
| Auditory Persistence | -6.1264 | 6.9559 | -0.88075 | 0.38259 |
| Visual Persistence | -1.1066 | 2.2277 | -0.49675 | 0.62154 |
| Auditory Sensory motor | 3.3571 | 5.565 | 0.60326 | 0.54896 |
